# Spontaneous depolarization-induced action potentials of ON-starburst amacrine cells during cholinergic and glutamatergic retinal waves

**DOI:** 10.1101/2020.10.21.349654

**Authors:** Rong-Shan Yan, Xiong-Li Yang, Yong-Mei Zhong, Dao-Qi Zhang

## Abstract

Correlated spontaneous activity in the developing retina (termed “retinal waves”) plays an instructive role in refining neural circuits of the visual system. Depolarizing (ON) and hyperpolarizing (OFF) starburst amacrine cells (SACs) initiate and propagate cholinergic retinal waves. Where cholinergic retinal waves stop, SACs are thought to be driven by glutamatergic retinal waves initiated by ON-bipolar cells. However, the properties and function of cholinergic and glutamatergic waves in ON- and OFF-SACs still remain poorly understood. As expected, we found that both SAC subtypes exhibited spontaneous rhythmic depolarization during cholinergic and glutamatergic waves. Interestingly, ON-SACs had wave-induced action potentials (APs) in an age-dependent manner, but OFF-SACs did not. We further found that the number of APs in ON-SACs was correlated with the amplitude of Ca^2+^ transients of either ON- or OFF-SACs during cholinergic retinal waves. These results advance the understanding of the cellular mechanisms underlying correlated spontaneous activity in the developing retina.

## 1. Introduction

Correlated spontaneous activity travels in a wave-like fashion (“retinal waves”) across the vertebrate retina before the emergence of vision [1,2]. Cholinergic retinal waves are initiated by and propagated between starburst amacrine cells (SACs), whereas glutamatergic retinal waves are generated by depolarizing (ON) bipolar cells [3–5]. Both types of waves are also transmitted vertically to retinal ganglion cells (RGCs), the output neurons of the retina, and then propagate along the optic nerve to the brain. Because they instruct the features of retinal projections, spatiotemporal patterns of retinal waves are critical for the establishment of eye-specific segregation and the formation of retinotopic maps [6–11].

SACs comprise mirror-symmetric ON and OFF subpopulations. ON-SACs have their somas in the ganglion cell layer (GCL) and dendrites stratified narrowly in the inner half of the inner plexiform layer (IPL), whereas OFF-SACs are located in the inner nuclear layer (INL), with their dendrites stratifying narrowly in the outer half of the IPL. ON-SACs are readily accessible and have been studied extensively in the developing retinas of rodents, including rabbits and mice [3,12–15]. They exhibit slow rhythmic spontaneous depolarization occasionally superimposed with small amplitudes of Ca^2+^ spikes during cholinergic waves [3,13,16]. Although positive current injection into rabbit ON-SACs can induce action potentials (APs) during development [17], it remains unknown whether APs are evoked by spontaneous slow depolarization of ON-SACs. APs mediate fast synaptic transmission; therefore, they could influence the initiation and propagation of cholinergic waves across the retina.

However, the cholinergic waves of OFF-SACs have not been characterized since these cells are located in the INL and are difficult to access. The density of OFF-SACs (~1200 cells/mm^2^) is slightly higher than that of ON-SACs (~1000 cells/mm^2^) [18,19]. Thus, OFF-SACs are thought to be as important as ON-SACs for the initiation, propagation, and termination of cholinergic waves. To understand the role of OFF-SACs, it is necessary to characterize the properties of cholinergic wave activity in them.

Cholinergic waves are replaced by glutamatergic retinal waves in rodents during the second postnatal week [14]. Glutamatergic waves propagate laterally between ON-bipolar cells and also travel to OFF-bipolar cells horizontally through inhibitory amacrine cells. Vertically, glutamatergic waves travel to RGCs and depolarize ON-RGCs and neighboring OFF-RGCs in sequence: ON before OFF [20,21]. Although SACs are important downstream neurons of bipolar cells, it remains unclear how ON- and OFF-SACs respond to input from bipolar cells during glutamatergic waves. This input could be important for understanding how glutamatergic waves regulate the development of SACs and the cholinergic system. In reverse, SACs could provide feedback to bipolar cells and feedforward to RGCs, which ultimately shapes signal transmission from bipolar cells to RGCs during glutamatergic waves.

In the present study, we used whole-cell patch-clamp recordings and Ca^2+^ imaging in combination with transgenic mice to systemically characterize the electrophysiological properties of genetically labeled ON- and OFF-SACs during cholinergic and glutamatergic waves. We find that ON-but not OFF-SACs exhibit wave-induced APs. We further find that APs enhance the Ca^2+^ intensities of ON- and OFF-SACs during cholinergic retinal waves.

## 2. Materials and Methods

### 2.1. Animals

Male and female neonate mice were used for the present study. Mice were housed in the Oakland University animal facility under a 12-h:12-h light-dark cycle, with access to food and water *ad libitum.* All experimental procedures were conducted according to the National Institutes of Health guidelines for laboratory animals and were approved by the Institutional Animal Care and Use Committees at Oakland University.

Choline acetyltransferase (ChAT) is the enzyme responsible for the synthesis of acetylcholine (ACh), and is specifically expressed in cholinergic neurons. Mice with ChAT(IRES)-Cre knock-in express Cre recombinase under the control of the ChAT promoter (stock No. 006410, Jackson Laboratory, Bar Harbor, ME). They were crossed with Ai9 mice in which a loxP-flanked STOP cassette prevents transcription of the red fluorescent protein variant tdTomato (stock No. 007909, Jackson Laboratory, Bar Harbor, ME) to produce ChAT-Cre/Ai9 (tdTomato) mice. After Cre exposure, this new mouse line expressed robust tdTomato fluorescence for visualizing SACs in the retina. In addition, enhanced green fluorescent protein (EGFP)–calmodulin–M13 fusion protein 6 fast variant (GCaMP6f) is a genetically encoded Ca^2+^ indicator. We crossed ChAT-Cre mice with Ai95D mice (stock No. 028865, Jackson Laboratory Bar Harbor, ME) in which a floxed STOP cassette prevents transcription of GCaMP6f to produce ChAT-Cre/Ai95D mice. After Cre exposure, ChAT-Cre/Ai95D mice express EGFP fluorescence in SACs following calcium binding. This mouse line allowed us to visualize SACs for electrophysiological recordings and Ca^2+^ imaging. We used ChAT-Cre/Ai9 and ChAT-Cre/Ai95D mice aged from postnatal day 3 (P3) to P14 (or before eye opening) for the present study.

### 2.2. Immunohistochemistry

Immunohistochemistry was performed in whole-mount retinas. Retinas were isolated and fixed with 4% paraformaldehyde (PFA). Fixed retinas were incubated overnight in blocking solution containing 1% bovine serum albumin (BSA) and 1% Triton X-200. After blocking, retinas were treated with primary antibodies for 2-3 days at 4 °C. Goat-anti-ChAT (1:500, Millipore Sigma-Aldrich, Burlington, MA) was paired either with rabbit-anti DsRed (1:1000, Invitrogen, Carlsbad, CA) for ChAT-Cre/Ai9 retinas or with rabbit anti-GFP (1:1000, Invitrogen) for ChAT-Cre/Ai95 retinas. Retinas were rinsed with 0.1 M PBS and then incubated with second antibodies at room temperature for 2 hours. Second antibodies included Alexa Fluor 488 donkey anti-goat IgG (1:500, Invitrogen), Alexa Fluor 594 goat anti-rabbit IgG (1:500, Invitrogen), Alexa Fluor 594 donkey anti-goat IgG (1:500, Invitrogen), and Alexa Fluor 488 goat anti-rabbit IgG (1:500, Invitrogen). Finally, retinas were rinsed and mounted with Vectashield (Vector Mounting Solution H-1400, Vector Laboratories, Burlingame, CA) for imaging. Images were taken using Zeiss Apotome 2 microscopy equipment (Zeiss, Oberkochen, Germany).

### 2.3. Patch-clamp electrophysiological recording

Pups were euthanized with an overdose of CO_2_ followed by cervical dislocation. Eyeballs were enucleated from euthanized pups and transferred to a dissection chamber, which contained extracellular solution bubbled with 95% O_2_ and 5% CO_2_ (pH ~7.4). Extracellular solution contained (in mM): 120 NaCl, 2.5 KCl, 1 MgSO_4_, 2 CaCl_2_, 1.25 NaH_2_PO_4_, 20 D-Glucose, 26 NaHCO_3_. Retinas were removed from eyeballs under dim red light. Each isolated retina was then transferred to and placed in a recording chamber with the ganglion cell side up. The recording chamber was mounted on the stage of an upright conventional fluorescence microscope (Axio Examiner, Zeiss, Oberkochen, Germany). Oxygenated extracellular solution was continuously perfused into the recording chamber at a rate of 2-3 ml/min. Solution temperature was maintained at 36-37 °C by a temperature control unit (TC-344A; Warner Instruments, Hamden, CT).

Prior to recording, retinas were kept in darkness for about 1 hour in order for them to adapt to the *ex vivo* environment. Retinas and recording pipettes were visualized on the computer monitor, which was connected to a digital camera (AxioCamMR3, Zeiss, Oberkochen, Germany) equipped on the microscope. GFP- and tdTomato-labeled cells in the retinas were identified with fluorescein isothiocyanate (FITC) and rhodamine filter sets, respectively, under fluorescence microscopy. Targeted cells and recording pipettes were monitored with infrared differential interference contrast (DIC) imaging for patch-clamp recordings except for simultaneous patch-clamp recording and Ca^2+^ imaging (see below). Whole-cell current-clamp recordings were obtained from SAC somata, and cell membrane potentials were amplified by an Axopatch 700B amplifier (Molecular Devices, San Jose, CA). If a recorded cell did not exhibit spontaneous APs, depolarizing current pulses of 0 to +250 pA (50 pA steps, 500 ms duration) were delivered into the cell to elicit AP discharge if the cell could generate them. Pipette resistance was 6~8 MΩ, with intracellular solution containing (in mM): 120 K-gluconate, 5 NaCl, 4 KCl, 10 HEPES, 2 EGTA, 4 Na_2_-ATP, 0.3 Mg_2_-GTP. Data were acquired using a Digidata 1440A digitizer and Clampex 10.7 software (Molecular Devices, San Jose, CA).

### 2.4. Calcium imaging

Spontaneous Ca^2+^ transients during retinal waves are reflected by changes in the intensity of GCaMP6f signals expressed in the retinas of ChAT-Cre/Ai95 mice. Fluorescence excited with a green light (470 ± 20 nm) was captured by an AxioCamMR3 camera and continuously visualized with AxioVision software. Real-time images were video recorded using the Open Broadcaster Studio software (bitrate: 2550 kbps, frame interval: 100 frames per second, output resolution: 1120 × 700 pixels). Images were acquired using a 63× or 20× water immersion objective lens.

### 2.5. Simultaneous Ca^2+^ imaging and patch-clamp recording

While the patch-clamp recording was conducted from SACs in ChAT-Cre/Ai95 retinas under the DIC, it was challenging to switch the DIC filter set to an FITC filter set for simultaneous Ca^2+^ imaging, as the switch usually physically disrupts the stability of the patch-clamp recording. To overcome this hurdle, we performed simultaneous patch-clamp recording and Ca^2+^ imaging under green fluorescent light with a FITC filter set. The tip of a glass pipette was coated with BSA-Alexa Fluor 488 conjugate (3 mM, Invitrogen). The coated pipette and an EGFP-labeled SAC were simultaneously visualized under green fluorescent light for patch-clamp recording. When the patch-clamp recording was stable, Ca^2+^ imaging was carried out without a filter set switch. If the patch-clamp recording was conducted in the GCL but Ca^2+^ imaging needed to be taken in the INL, we gently refocused the imaging plane from the GCL to the INL.

### 2.6. Drugs

The sodium channel blocker tetrodotoxin (TTX), nicotinic ACh receptor antagonists (hexamethonium bromide and DhβE), and glutamate receptor antagonists (NBQX or CNQX for AMPA receptors and D-AP5 for NMDA receptors) were used for electrophysiological recording and calcium imaging experiments. Chemicals were obtained from Tocris Bioscience (Ellisville, MO, USA), Sigma-RBI (or MilliporeSigma, St. Louis, MO, USA), and Hello Bio Inc. (Princeton, NJ, USA). Drugs were stored in frozen stock solutions and dissolved in extracellular medium before an experiment.

### 2.7. Data analysis

The resting membrane potentials (RMP) and spontaneous depolarization amplitudes of SACs, as well as the frequency of APs superposed on a depolarization, were measured offline using Clampfit 10.7 (Molecular Devices, Sunnyvale, CA). The number of spontaneous depolarization events in each SAC was counted manually over a recording time (min). The frequency (F) of the spontaneous depolarization was determined by dividing the number of events by the recording time (events/min). Raw recording traces were exported using Clampfit 10.7 and were edited for presentation using CorelDRAW Home & Student 2019 (Professional graphic design software).

Ca^2+^ images were extracted and converted into GIF format offline with video editing software BeeCut. The export settings were 1280 × 720 pixel resolution, 12 frames/s speed, and 6000 bps bitrate. Exported images were further analyzed using the plug-in *Time Series Analyzer* (Ver. 3) in Fiji ImageJ (LOCI, University of Wisconsin-Madison).

For analysis of the simultaneous Ca^2+^ imaging and patch-clamp recordings, we selected a circular region of interest (ROI) that included a recorded SAC and measured intensities of GCaMP6f signal within the ROI from a series of images taken over time. We plotted the intensities (arbitrary unit) as a function of recording time to construct the Ca^2+^ wave trace.

We then determined the relationship between APs of ON-SACs and the Ca^2+^ waves in the INL or GCL. For each simultaneous recording, we obtained the adjusted peak amplitude of individual Ca^2+^ waves by subtracting the mean baseline Ca^2+^ intensity value before the wave from the actual peak value of Ca^2+^ intensity during the wave. The adjusted peak amplitude of each wave was normalized by dividing it by the maximum peak amplitude observed during the entire recording. For every Ca^2+^ wave, we counted the number of APs from its corresponding recorded ON-SAC electrical wave. We plotted the normalized peak amplitudes of Ca^2+^ waves as a function of the number of APs of ON-SACs. The data points were fit with a linear regression equation, yielding the coefficient of determination (*r*^2^) and probability (*p*) value for the coefficient, using Sigma Plot 14.0 (San Jose, California). A value of *p* < 0.001 was considered significantly correlated.

Two independent groups with normally distributed data but unequal sample sizes were compared using a *t*-test in Prism 8.0 GraphPad (San Diego, CA); similarly, when the distribution was not normal, a nonparametric t-test with Welch’s correction was employed. A paired *t*-test was used to compare the means of the two related groups. Values were presented as the mean ± standard error of mean (SEM). A value of *p* < 0.05 was considered statistically significant.

## 3. Results

### 3.1. Characterization of genetically labeled SACs in transgenic mouse lines

To visualize SACs for electrophysiological recordings, we produced ChAT-Cre/tdTomato (Ai9) transgenic mice in which SACs expressed a tdTomato fluorescent protein (see Methods). td-Tomato-expressing cells were imaged from living retinas beginning at P3 under fluorescent microscopy. We found that they were distributed in the GCL as well as in the INL (Figure 1A). To confirm whether tdTomato-expressing cells were SACs, P5 retinas were immunostained with antibodies against ChAT and DsRed, respectively (Figure 1B). We found that all DsRed-expressing cells were ChAT positive (ChAT^+^) ON-SACs or OFF-SACs, indicating that tdTomato specifically marked SACs in this transgenic mouse line. Although we occasionally observed ChAT^+^ SACs without DsRed staining (Figure 1B, arrows), this mouse line is a robust animal model for use in visualizing living SACs for patch-clamp recordings.

**Figure 1.**
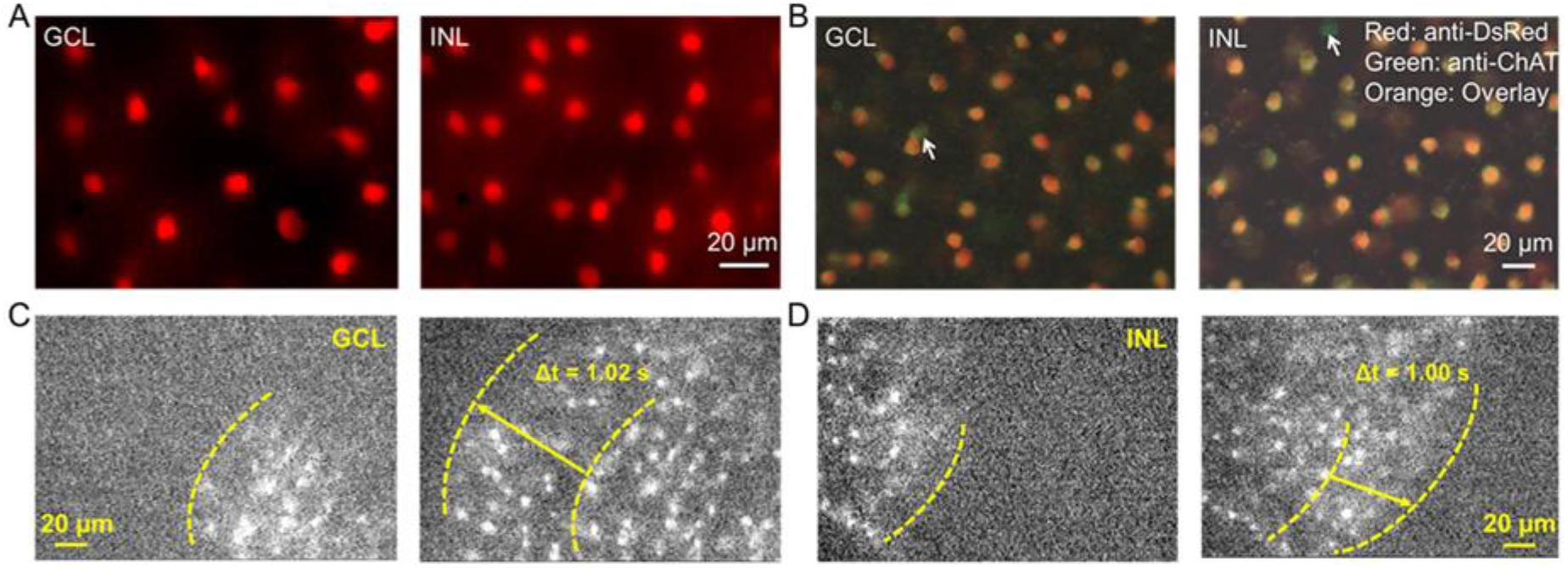
Genetically labeled ON- and OFF-SACs in mouse retina. (A) Images of a living retina isolated from a ChAT-Cre/tdTomato mouse at P5, with tdTomato-labeled SACs (red) in the ganglion cell layer (GCL, left) and inner nuclear layer (INL, right). (B) Merged images of a fixed retina from a ChAT-Cre/tdTomato mouse at P5 showing colocalizations (orange) of immunostaining with anti-DsRed for tdTomato-marked cells (red) and anti-ChAT for SACs (green) in the GCL (ON-SAC, left) and INL (OFF-SACs, right). In both layers, the DsRed-labeled cells all tested positive for the ChAT antibody. The arrows point to the ChAT positive cells that were not labeled with the DsRed antibody. (C) Two consecutive images taken in the GCL from a living retina of a ChAT-Cre/GCaMP6f mouse at P5 at a 1.02-s interval (Δt) showed that a Ca^2+^ wave traveled across ON-SACs from the lower-right corner (left panel) to the upper-left corner (right panel). (D) Two consecutive images taken in the INL from a living retina of a *ChAT-Cre/*GCaMP6f mouse at P5 with a 1-s interval (Δt) showed that a Ca^2+^ wave moved across OFF-SACs from the upper-left corner (left panel) to the lower-right corner (right panel). (C) and (D): *The* yellow dashed lines indicate the Ca^2+^ wave front. The arrows indicate the direction of the Ca^2+^ wave’s propagation.

To monitor Ca^2+^ transients across developing SACs, we replaced the tdTomato reporter with a GCaMP6f reporter for ChAT-Cre mice, producing ChAT-Cre/GCaMP6f (Ai95D) mice (see Methods). In this mouse line, we observed EGFP-expressing cells in living retinas (Figures 1C and D). These cells were immunoreactive to antibodies against ChAT and GFP (data not shown), indicating that EGFP specifically labels ON- and OFF-SACs. We monitored Ca^2+^ waves of SACs by continually imaging *GCaMP6f* expression over time in the GCL (Figure 1C) and INL (Figure 1D), respectively. In Figure 1C, a Ca^2+^ wave of ON-SACs traveled from the lower-right corner (left panel) to the upper-left corner (right panel). OFF-SACs also exhibited Ca^2+^ waves. One example is illustrated in Figure 1D, demonstrating that a wave was propagated from the upper-left corner (left panel) to the lower-right corner (right panel). These results suggest that ChAT-Cre/Ai95D mice can be used to monitor the Ca^2+^ waves of SACs as well as to visualize SACs for patch-clamp recordings.

### 3.2. ON-SACs exhibit spontaneous depolarization accompanied by APs during cholinergic waves

To determine the physiological properties of ON-SACs in the developing mouse retina, we performed whole-cell current-clamp recordings from tdTomato-expressing ON-SACs in flat-mount retinas. The advantage of this retina preparation is that the neural networks of SACs remain intact, which is required for the wave initiation and propagation [4,13]. Since mouse cholinergic retinal waves appear in the first postnatal week [4,13], we began recordings from retinas at P5. As expected, we found that all ON-SACs recorded (*n* = 28) depolarized spontaneously from the RMP with a slow onset (Figure 2A and 2B). However, we unexpectedly found that in 17 out of 28 cells (61%), each slow depolarization was accompanied by a burst of APs with a rate of 3.4 ~ 10 Hz (Figure 2A and 2D). This percentage (61%) at P5 was increased to 78% at P3–P4, but dropped to 21% at P6–P7 (Figure 2D), indicating the number of ON-SACs with spontaneous depolarization-induced APs is determined in an age-dependent manner. ON-SACs with and without APs are hereafter referred to as spontaneously spiking (S-spiking) and non-spontaneously spiking (non-S-spiking) ON-SACs, respectively. The mean frequency of spontaneous depolarization between S-spiking (0.73 ± 0.45 events/min, *n* = 31) and non-S-spiking ON-SACs (0.67 ± 0.03 events/min, *n* = 40) had no significant difference (Figure 2E; *p* = 0.27).

**Figure 2.**
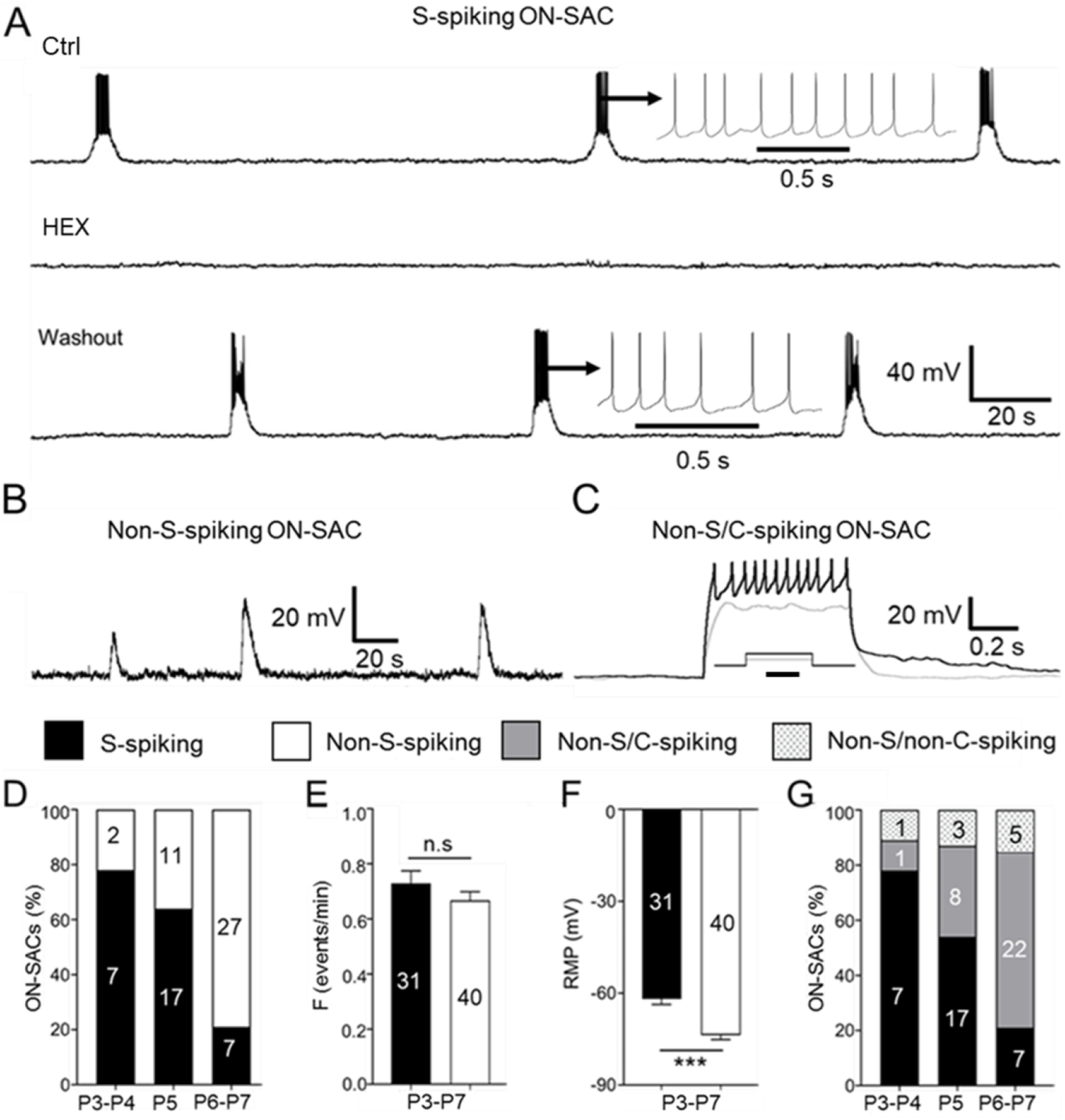
Spontaneous depolarization of ON-SACs accompanied by a burst of APs during cholinergic retinal waves. Whole-cell current-clamp recordings were performed on ON-SACs from flat-mount retinas of ChAT-Cre/tdTomato mice at P3–P7. (A) An ON-SAC recorded at P5 exhibited spontaneous rhythmic depolarization, and each of depolarization was accompanied by a burst of APs (top trace). An insert highlights the burst of APs on an extended time scale. The spontaneous depolarization and APs were blocked by HEX (100 μM) (middle trace), and the blockage was reversible upon washout (bottom trace). (B) An ON-SAC exhibited spontaneous rhythmic depolarization without APs at P5. (C) An ON-SAC did not have spontaneous depolarization-induced APs at P5 but showed current injection–induced APs with reduced amplitudes (black trace). An insert shows the injected currents (gray: 200 pA, black: 250 pA) at 500 ms (scale bar = 250 ms). Therefore, the ON-SACs had spontaneously spiking (S-spiking) and non-S-spiking subtypes. The latter is further classified into non-S-spiking with (non-S/C-spiking) and without current-induced APs (non-S/non-C-spiking). The S-spiking, non-S-spiking, non-S/C-spiking, and non-S/non-C-spiking ON-SACs are presented as black, white, and gray bars and as black-dotted bars on a white background, respectively, in (D) through (G). (D) The percentages of S-spiking and non-S-spiking ON-SACs at P3–P4, P5, and P6–P7, respectively. (E) The mean data pooled from P3–P7 showed no significant difference (n.s.) in the spontaneous depolarization frequency (events/min) between the S-spiking and non-S-spiking ON-SACs (p = 0.487). (F) The mean data pooled from P3–P7 demonstrated that the resting membrane potential (RMP) for S-spiking ON-SACs was significantly higher than that of non-S-spiking ON-SACs (***p < 0.001). (G) The percentages of S-spiking, non-S/C-spiking, and non-S/non-C-spiking ON-SACs at P3–P4, P5, and P6–P7, respectively. The numbers within the bars in (D) through (G) indicate the numbers of ON-SACs tested. The means ± SEM are presented in (D) through (G).

To confirm whether the spontaneous depolarizations were mediated by ACh receptors, we bath-applied ACh receptor antagonist hexamethonium (HEX, 100 μM) or DHβE (3 μM) to the retinas. We found that both HEX (Figure 2A) and DhβE (data not shown) eliminated spontaneous depolarizations and their associated APs in S-spiking ON-SACs. HEX also completely blocked the spontaneous depolarization of non-S-spiking ON-SACs (data not shown, *n* = 18).

Further analysis showed that the mean RMP of S-spiking ON-SACs (−61.90 ± 1.80 mV, *n* = 31) was significantly more positive than that of non-S-spiking ON-SACs (−73.72 ± 1.45, *n* = 40; Figure 2F; *p* < 0.001). This 12-mV membrane potential hyperpolarization could keep non-S-spiking ON-SACs from reaching the threshold potential for AP generation. However, we found that the spontaneous depolarization peak amplitude of non-S-spiking ON-SACs (31.68 ± 1.94 mV, *n* = 40) was approximately 9 mV higher than that of S-spiking ON-SACs (22.77 ± 1.29 mV, *n* = 31; *p* < 0.001). Although this 9-mV depolarization almost canceled out the 12-mV hyperpolarization of the RMP in non-S-spiking ON-SACs, they still could not fire APs. The results imply that non-S-spiking ON-SACs either have a higher threshold for AP generation or do not have an ability to generate APs.

To distinguish these two possibilities, we injected positive currents into non-S-spiking ON-SACs to determine whether positive currents can induce APs. Eight of the 11 cells (73%) exhibited current-induced APs but with reduced amplitude at P5 (Figure 2C and 2G). Similar results were obtained in 81% of non-S-spiking ON-SACs at P6-7 (Figure 2G). This result indicates that the majority of non-S-spiking ON-SACs can generate APs if their membrane potentials depolarize enough to reach the AP threshold membrane potential. We referred to this subtype as non-S/C-spiking ON-SACs. However, the rest of the non-S-spiking ON-SACs at P5 or P6–P7 did not have current-induced APs (Figure 2G); these were referred to as non-S/non-C-spiking ON-SACs, indicating that a minority of ON-SACs do not have the ability to generate APs.

### 3.3. OFF-SACs show spontaneous depolarization without APs during cholinergic waves

Research to date has not uncovered the electrical wave properties of OFF-SACs in developing retinas. Here, we successfully patched OFF-SACs from the INL in a flat-mount retina. All OFF-SACs recorded from P3 to P7 exhibited spontaneous slow rhythmic depolarization, but these depolarizations did not induce APs (Figure 3A, top trace). The spontaneous depolarizations of OFF-SACs were completely blocked by HEX (Figure 3A, bottom trace) or DhβE (data not shown), suggesting that they were mediated by cholinergic transmission (*n* = 8). The frequency of spontaneous depolarization was 0.55 ± 0.03 events/min (*n* = 21), which was significantly lower than that of S-spiking (0.73 ± 0.05 events/min, *n* = 31, *p* < 0.05) or non-S-spiking ON-SACs (0.67 ± 0.03 events/min, *n* = 40, *p* < 0.05). The mean RMP of OFF-SACs was −77.29 ± 2.46 mV, which was similar to that of non-S-spiking ON-SACs (−73.72 ± 1.45 mV, *p* = 0.28) but significantly lower than that of S-spiking ON-SACs (−61.90 ± 1.80 mV, *p* < 0.001). However, the peak amplitude of OFF-SAC spontaneous depolarization (20.79 ± 2.00 mV) was similar to that of S-spiking ON-SACs (22.77 ± 1.29 mV, *p* = 0.37) and significantly smaller than that of non-S-Spiking ON-SACs (31.68 ± 1.94 mV, *p* < 0.001). Positive current-induced APs were only observed in 15% of OFF-SACs (non-S/C-spiking OFF-SACs, Figure 3B and 3D). A majority of these cells (85%) showed no current-induced APs (non-S/non-C-spiking OFF-SACs, Figure 3C and 3D). These results indicate that OFF-SACs have much less ability to generate APs than ON-SACs, even if spontaneous depolarizations reach a threshold level for AP generation

**Figure 3.**
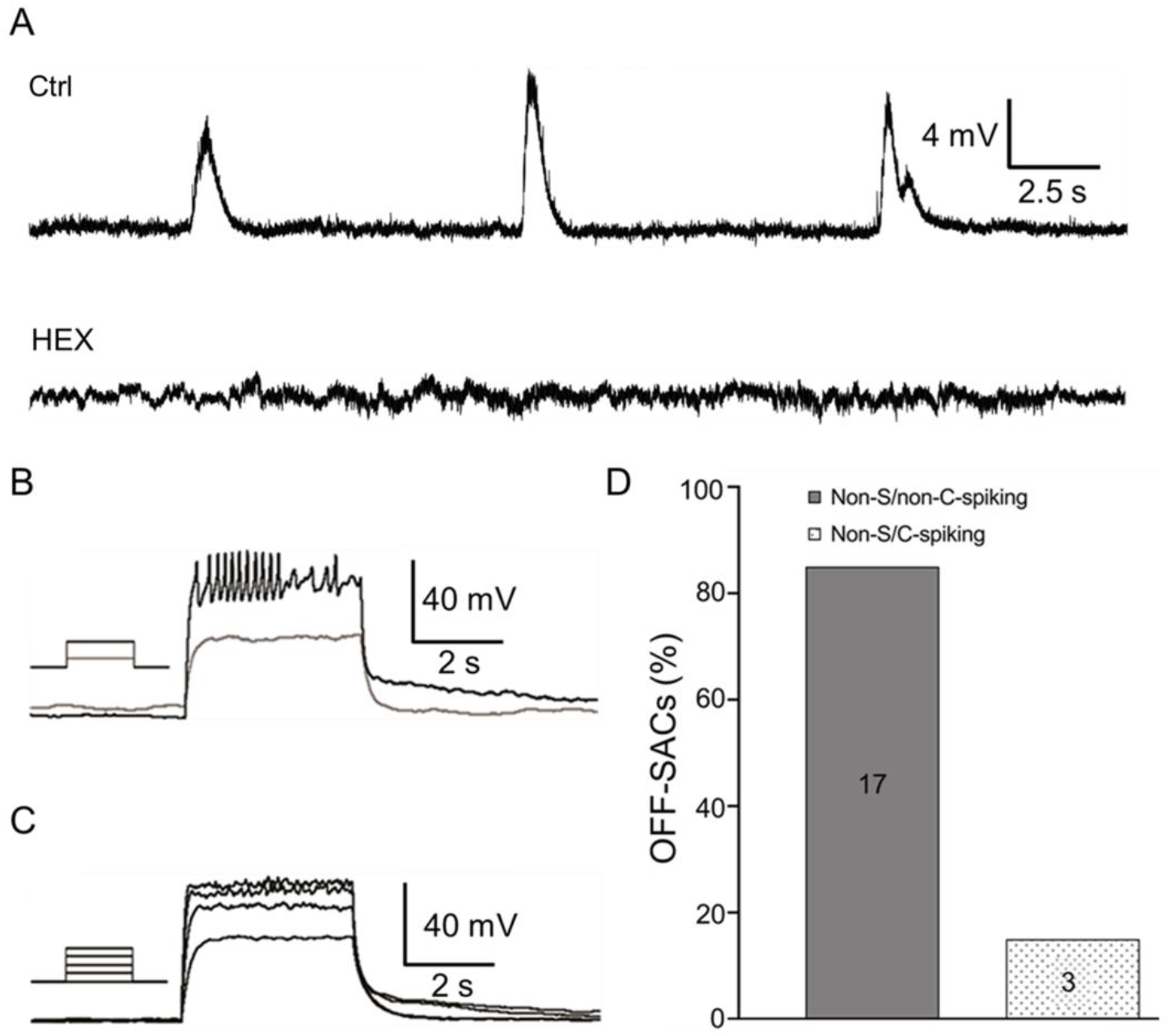
OFF-SACs exhibit spontaneous rhythmic depolarization without APs. Whole-cell current-clamp recordings were conducted on OFF-SACs from flat-mount retinas of ChAT-Cre/tdTomato mice at P3–P7. (A) An OFF-SAC was recorded at P5. The cell exhibited spontaneous rhythmic depolarization (top trace). HEX (100 μM) completely blocked the spontaneous activity (bottom trace). (B) An OFF-SAC showed spontaneous depolarization at P3. When 150 pA and 200 pA positive currents of 500 ms (see the inserted protocol) were injected, the latter induced depolarization, which was accompanied by APs with reduced amplitudes. (C) An OFF-SAC showed spontaneous depolarization at P4. A series of positive currents of 500 ms from 50 pA to 200 pA, at 50 pA steps, were injected into the cell. The currents induced depolarization without having APs. (D) The data collected from P3 to P7 were pooled. The bars show the percentages of OFF-SACs with (non-S/C-spiking) and without (non-S/non-C-spiking) current-induced APs. The number of OFF-SACs tested is indicated within the bars.

### 3.4. S-spiking SACs still only appear in the ON layer during glutamatergic retinal waves

Glutamatergic retinal waves initiated by ON-bipolar cells appear in the second postnatal week [20]. To determine whether this type of retinal wave drives ON- and OFF-SACs, we conducted whole-cell current-clamp recordings on them from P8 to P14 and found that both ON- and OFF-SACs recorded exhibited spontaneous depolarization (Figure 4A and 4B, top traces). Interestingly, spontaneous depolarization of both subtypes had two peaks; the second peak often appeared before the first peak returned to the baseline (Figure 4A and 4B, top traces). In addition, 26% of ON-SACs (5 of 19) exhibited spontaneous depolarization-induced bursts of APs at P8–P9 (Figure 4C), but this percentage dropped to 17% (2 of 12) at P10–P14, further indicating that S-spiking ON-SACs are age-dependent. None of the OFF-SACs recorded showed spontaneous depolarization-induced APs. The mean spontaneous depolarization frequency of OFF-SACs (1.41 ± 0.23 events/min, *n* = 13) was higher than that of ON-SACs with APs (0.77 ± 0.17 events/min, *n* = 8, *p* < 0.05) and that of non-S-spiking ON-SACs (1.04 ± 0.15 events/min, *n* = 26, but *p* = 0.186), respectively.

**Figure 4.**
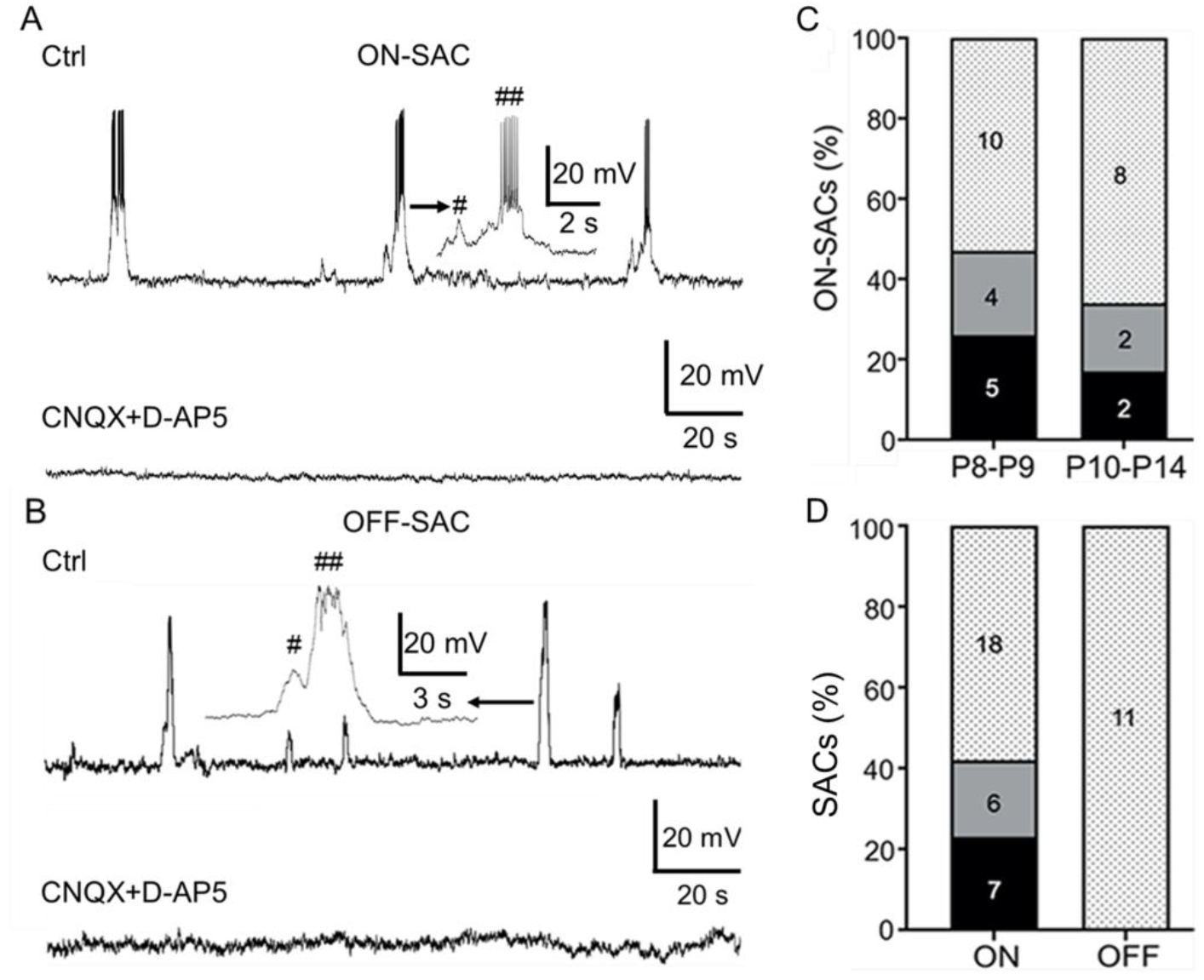
Spontaneous activity of ON- and OFF-SACs during glutamatergic waves. ON- and OFF-SACs from flat-mount retinas of ChAT-Cre/tdTomato mice were recorded from P8 to P14 using a whole-cell current-clamp configuration. (A) An ON-SAC exhibited spontaneous rhythmic depolarization, and most of the depolarization had two peaks at P9 (top trace). An insert illustrates that the APs were superimposed onto the initial (#) and delayed peaks (##) on an extended time scale. CNQX (50 μM) and D-AP5 (50 μM) completely blocked the spontaneous activity (bottom trace). (B) An OFF-SAC also exhibited spontaneous rhythmic depolarization at P9, and the depolarization had initial (#) and delayed (##) peaks (see the insert with an extended time scale). CNQX (50 μM) and D-AP5 (50 μM) completely blocked the spontaneous activity (bottom trace). (C) The ON-SACs were pooled into two groups: P8–P9 and P10–P14. The bars show the percentages of S-spiking (black), non-S/C-spiking (gray), and non-S/non-C-spiking (black dots) ON- and OFF-SACs. (D) The ON- and OFF-SACs recorded from P8–P14 were pooled. The bars illustrate the percentages of S-spiking (black), non-S/C-spiking (gray), and non-S/non-C-spiking ON-SACs (block dots) for each SAC subtype. The number of SACs tested is presented within the bars for (C) and (D).

To confirm that the depolarizations were mediated by glutamate receptors, we applied glutamate receptor antagonists CNQX (or NBQX, 50 μM) and D-AP5 (50 μM). We found these receptor antagonists completely blocked spontaneous depolarization of ON-(Figure 4A, bottom trace, *n* = 22) and OFF-SACs (Figure 4B, bottom trace, *n* = 7).

To determine whether non-S-spiking ON- and OFF-SACs are able to generate APs, we injected positive currents into these cells. We found that the current injection induced APs in 21% of ON-SACs (4 of 19) at P8–P9 and 17% (2 of 12) at P10–P14 (Figure 4C). When we pooled all ON-SACs recorded from P8–P14 together, 23% of ON-SACs exhibited spontaneous depolarization-induced APs, 19% had no S-spiking but with current-induced APs, and 58% did not have any ability to generate APs (Figure 4D). In contrast, none of the OFF-SACs recorded (0 of 11) exhibited current-induced APs from P8–P14 (Figure 4D). Our data suggest that some ON-SACs are still capable of generating APs during glutamatergic waves, but OFF-SACs are not.

### 3.5. Blockage of spontaneous and current-induced APs of SACs by TTX

Next, we determined whether sodium channels mediated spontaneous and current-induced APs in SACs. We found that TTX, a specific sodium channel blocker, eliminated the spontaneous APs during cholinergic waves (Figure 5A). We further found that the wave frequency was not significantly changed in the presence of TTX (Figure 5B, right panel, *p* = 0.38, *n* = 10). However, the peak amplitude of spontaneous depolarization was significantly augmented after the blockage of APs by TTX (Figure 5B, left panel, *p* < 0.05, *n* = 10). The data suggest that ACh-mediated depolarization does not reach its peak while APs are induced. TTX also blocked current-induced APs in ON-SACs (Figure 5C, *n* = 13), as well as in OFF-SACs (Figure 5D, *n* = 3). The data indicate that both spontaneous depolarization and current-induced APs of SACs are mediated by TTX-sensitive sodium channels.

**Figure 5.**
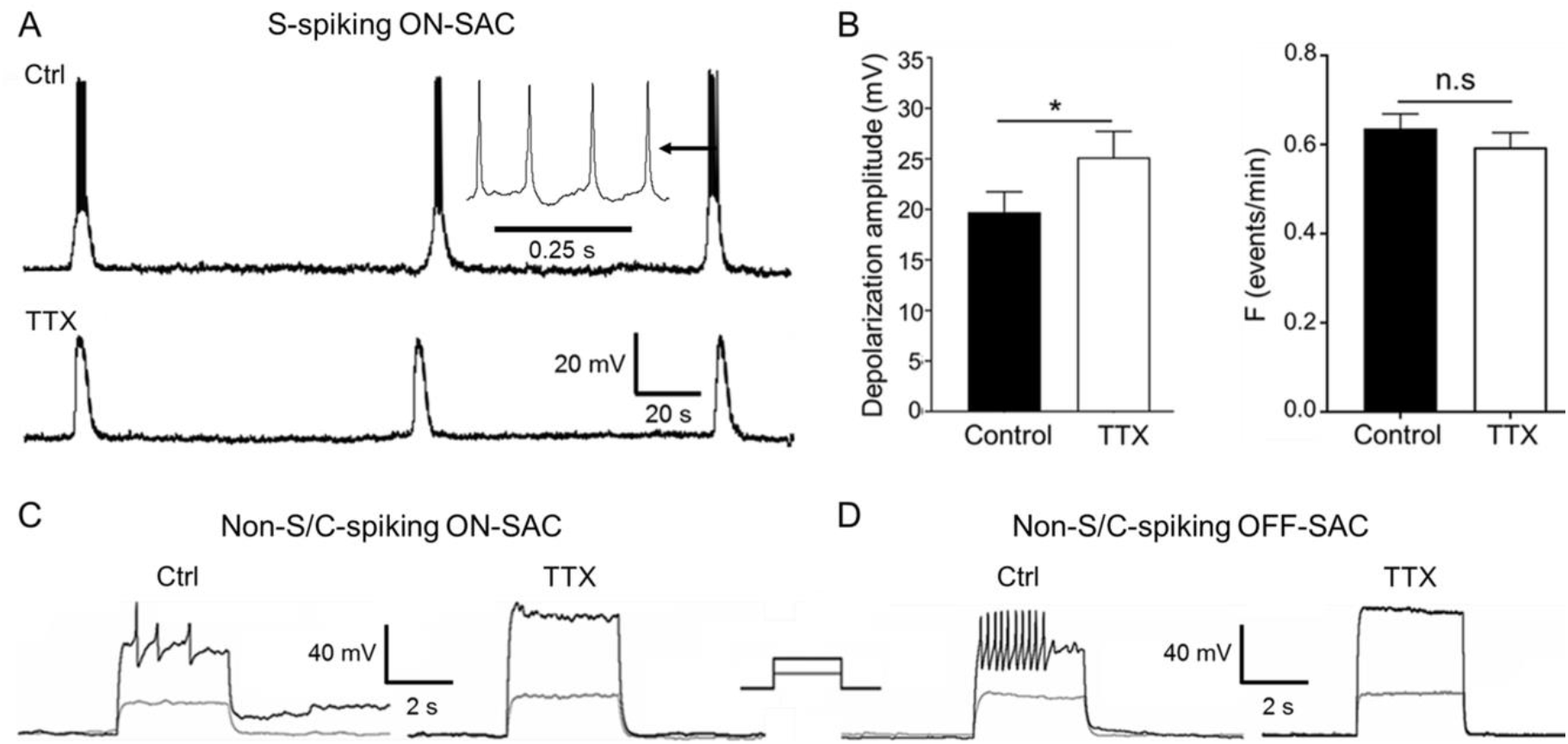
TTX blocking spontaneous APs and current-induced spikes in SACs. ON- and OFF-SACs from flat-mount retinas of ChAT-Cre/tdTomato mice were recorded from P3 to P7 using a whole-cell current-clamp configuration. (A) APs (see the insert) were superposed onto an S-spiking ON-SAC experiencing spontaneous depolarization in the control (top trace), but they were eliminated in the presence of 0.3 μM TTX (bottom trace) at P5. (B) 0.1-0.5 μM TTX blocked spontaneous depolarization-induced APs in S-spiking ON-SACs recorded from P3 to P7. Pooled data demonstrated that TTX significantly increased the peak amplitude of spontaneous depolarization (left panel, n = 10, *p < 0.05, mean ± SEM) but did not significantly change (n.s.) the frequency of spontaneous depolarization (right panel, n = 10, p = 0.38, mean ± SEM). (C) In a non-S-spiking ON-SAC at P5, current-induced spikes (left panel) were blocked by 0.3 μM TTX (right panel). (D) In a non-S-spiking OFF-SAC at P4, current-induced spikes (left panel) were blocked by 0.3 μM TTX (right panel). The insert for (C) and (D) illustrates the injected currents (gray: 200 pA, black: 250 pA) of 500 ms.

### 3.6. Relationships between APs of ON-SACs and Ca^2+^ waves in the GCL and INL

Our data and previous reports clearly demonstrate that ACh released from SACs during cholinergic waves or glutamate released from bipolar cells during glutamatergic waves produces postsynaptic depolarization in ON- and OFF-SACs [13,14]. The depolarization of some ON-SACs induces TTX-sensitive APs (Figures 2 and 4). Both graded potentials and APs can activate voltage-dependent Ca^2+^ channels, increasing intracellular Ca^2+^ during a wave. To characterize the relationship between APs and Ca^2+^ transients of ON-SACs during cholinergic waves, we performed simultaneous electrical recordings and Ca^2+^ imaging from ON-SACs in the GCL. Due to the dilution of intracellular contents, we did not construct the changes in calcium intensity from the patched SAC. Instead, we referred to the recorded SAC as the center of a circular ROI (size: 2996.2 μm^2^), which included any recorded S-spiking ON-SAC and 3 to 4 of its neighboring cells. Such ROIs were used for Ca^2+^ signal analysis in the GCL. The advantage of this analysis is that changes in the intensity of these regions also reflect intracellular Ca^2+^ transients from neighboring SACs, which could be driven by a patched S-spiking SAC. We found that each electrical wave of the S-spiking SAC (Figure 6A1 and 6A1’) was followed with a Ca^2+^ wave measured from the ROI in the GCL (Figure 6A2 and 6A2’).

**Figure 6.**
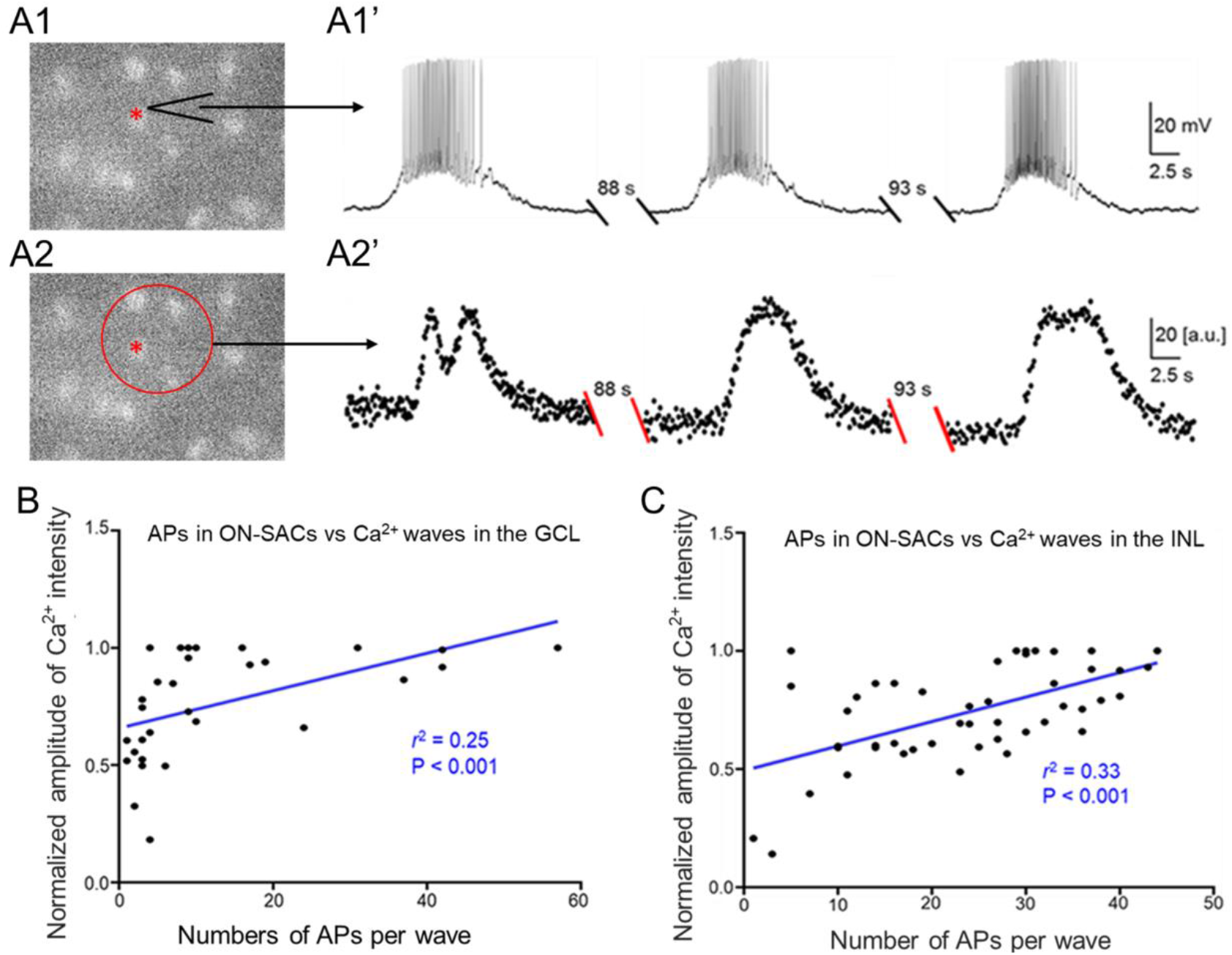
The APs of ON-SACs influencing the Ca^2+^ intensities of the GCL and INL during cholinergic waves. (A) Simultaneous Ca^2+^ imaging and whole-cell current-clamp recordings were performed in the GCL of retinas isolated from ChAT-Cre/Ai95 mice at P4–P5. A live image in A1 shows GCaMP6f-labeled ON-SACs at P5. One of the ON-SACs (indicated by a red star) was patched with a glass pipette (indicated by an arrow). The cell exhibited spontaneous rhythmic depolarization accompanied by APs (A1’). Ca^2+^ images in the GCL were taken simultaneously with the ON-SAC recording. An ROI (a red circle in A2) was selected to construct a Ca^2+^ wave trace. The trace demonstrates that each Ca^2+^ transient was preceded by an electrical wave from the ON-SAC (A2’). In A1’ and A2’, the recording trace between each wave was not shown (double slashes) for a period of time (indicated above the double-slashes). (B) The relationship was plotted (black dots) between the normalized peak amplitude of a Ca^2+^ wave in the GCL and the number of APs in the corresponding ON-SAC electrical wave. The scatterplot data were fit with the linear regression (blue line, r^2^ = 0.25, p < 0.001). (C) Ca^2+^ imaging in the INL was simultaneously performed alongside whole-cell current-clamp recordings of ON-SACs in the GCL in retinas from ChAT-Cre/Ai95 mice at P4–P5. The normalized peak amplitude of a Ca^2+^ wave in the INL was plotted as a function of the number of APs of the corresponding ON-SAC electrical wave (black dots). The group data were fit with the linear regression (blue line, r^2^ = 0.33, p < 0.001).

To determine whether APs in SACs contribute to the strength of Ca^2+^ waves, we counted the number of APs from each electrical wave of S-spiking SACs and measured the peak intensity of the Ca^2+^ wave of the ROI in the GCL. We plotted the normalized peak Ca^2+^ intensity (see Methods) as a function of the number of APs superimposed on the corresponding spontaneous depolarization of ON-SACs (Figure 6B). The data were fit with a linear regression, yielding a significant coefficient of determination (*r*^2^ = 0.25, *p* < 0.001, Figure 6B). This result suggests that APs of ON-SACs increase the Ca^2+^ transients across the GCL during cholinergic waves.

To determine whether APs of ON-SACs also influence Ca^2+^ waves in the INL, we performed electrophysiological recordings from S-spiking ON-SACs and simultaneously imaged the Ca^2+^ transients of OFF-SACs across the INL using P4 and P5 ChAT-Cre/Ai95D mice. A circular ROI (2996.2 μm^2^) of the INL located vertically above patched ON-SACs was used to construct changes in Ca^2+^ transients over time. We counted the number of APs from each electrical wave of S-spiking ON-SACs, measured the peak intensity of the corresponding Ca^2+^ waves of the ROI in the INL, and constructed a scatterplot between the number of APs and the normalized peak Ca^2+^ intensity (Figure 6C). The scatterplot data were fit with linear regression, yielding a significant coefficient of determination (*r*^2^ = 0.33, *p* < 0.001, Figure 6C). This result suggests that APs of ON-SACs also influence Ca^2+^ transients across the INL during cholinergic waves.

## 4. Discussion

In the present study, our data demonstrate that ON- and OFF-SACs exhibited spontaneous rhythmic depolarization during cholinergic and glutamatergic waves, respectively. The spontaneous depolarizations of ON-SACs were accompanied by TTX-sensitive APs in an age-dependent manner. However, APs are not induced by the depolarization of OFF-SACs. We further found that the APs of ON-SACs enhanced the intensity of Ca^2+^ waves across the GCL and INL. Our results suggest that developing ON-SACs can generate APs that promote the strength of Ca^2+^ waves across the retina.

### 4.1. Distinct electrical properties of ON- and OFF-SACs during cholinergic waves

Our results demonstrate that cholinergic waves of ON-SACs consist of two components: ACh-induced graded depolarization and a depolarization-induced, sodium channel-mediated AP. The latter component disagrees with previous reports showing that ACh-induced graded depolarizations in both rabbit and mouse ON-SACs are accompanied by repetitive Ca^2+^ spikes [3,13,16]. Although the cause of this inconsistency is unclear, the ChAT-Cre knock-in and fluorescence protein reporters are unlikely to contribute because not all ON-SACs can fire APs. Notably, our results are supported by the previously reported current injection-induced APs of ON-SACs in rabbit vertical retinal slice preparations [17]. Taken together, the results indicate that ON-SACs contain sodium channel proteins that mediate APs.

Based on their ability to generate APs, ON-SACs can be classified into S-spiking, non-S/C-spiking, and non-S/non-C-spiking subtypes. This classification could be determined by three parameters: the RMP, the amplitude of depolarization, and the density of the sodium channel’s protein expression. We found that the RMP of non-S-spiking ON-SACs were more negative than those of S-spiking ON-SACs were, which could keep them from reaching the threshold potential for sodium channel activation. However, this is unlikely to be the case because the depolarization amplitude of non-S-spiking ON-SACs was significantly larger than that of S-spiking ON-SACs. When positive currents were injected into non-S-spiking ON-SACs, some of the cells exhibited APs with reduced amplitude. However, a small percentage of them did not show any current-induced spikes, indicating that these SACs may express no or fewer sodium channel proteins with which to generate APs. We further found that the percentage of S-spiking ON-SACs decreased as postnatal days increased, indicating that the expression of sodium channel proteins in ON-SACs is significantly reduced in an age-dependent manner.

In contrast, OFF-SACs showed no spontaneous depolarization-induced APs during cholinergic waves. A majority of them did not even have current-induced spikes, suggesting that OFF-SACs have less or no sodium channel protein expression with which to mediate APs. Both ON- and OFF-SACs are born during an embryonic stage of development [22]. It is unknown whether sodium channel proteins are produced in both subtypes of SACs at birth. If they are, then our data suggest that the expression of sodium channels in OFF-SACs disappears earlier than in ON-SACs. Alternatively, our interpretation would be that OFF-SACs produce less sodium channel proteins than ON-SACs do from birth onward. Further investigation is needed to validate these electrophysiological data with specific antibodies against subunits of sodium channel proteins in both SAC subtypes during development.

### 4.2. Properties and mechanisms of glutamatergic waves in ON- and OFF-SACs

Here, we demonstrate for the first time that both ON- and OFF-SACs exhibit spontaneous rhythmic depolarization in the second postnatal week, indicating that they are driven by glutamatergic waves generated in ON-bipolar cells. Interestingly, the depolarization from both subtypes had two peaks, indicating that ON- and OFF-SACs may receive inputs from ON- and OFF-bipolar cells, respectively. Our interpretation is that ON-bipolar cells spontaneously release glutamate onto ON-SACs via a synaptic mechanism in the ON layer of the retina, producing the first peak of the wave. Glutamate may spill over extrasynaptically and diffuse to the OFF layer, thus generating the first peak of the wave in OFF-SACs [23]. In addition, the glutamate released from ON-bipolar cells activates inhibitory amacrine cells, which send an inhibition to OFF bipolar cells [20,21]. When glutamate release stops and the inhibition of OFF bipolar cells is relieved, OFF bipolar cells can rebound from the hyperpolarization. This rebound depolarization can stimulate glutamate release in the OFF layer [20,21]. Glutamate likely diffuses into the ON and OFF layers, thus producing the second peak of the wave in ON- and OFF-SACs.

Notably, ON-SACs can generate APs during glutamatergic waves, but OFF-SACs do not. The percentage of ON-SACs that can generate APs further dropped compared to that during the period of cholinergic waves. The data again suggest that the expression of sodium channels on ON-SACs could decrease in an age-dependent manner. The existence of TTX-sensitive sodium channels in SACs from mature retinas had been disputed [17,24–30]. In particular, a recent systematic study did not find evidence for TTX-sensitive sodium channels [30]. However, mature SACs appear to express TTX-resistant sodium channels, which play a role in direction selectivity [31–33]. However, our data do not support that TTX-resistant sodium channels are expressed in developing SACs (Figure 5).

In the GCL, ON-SACs and RGCs represent 11.5% and 41% of the total number of neurons, respectively [34]. When a method of microelectrode array (MEA) is used to evaluate cholinergic and glutamatergic waves from neurons in the GCL, both spiking SACs and RGCs can be recorded. Unfortunately, MEA extracellular recordings cannot distinguish spiking ON-SACs from RGCs. Since ~20% of recordings could be from SACs, they should be considered when interpreting MEA data from the developing retina.

### 4.3. *Possible roles of ON-SAC’s APs in the early* development of the visual system

Our data show the correlation between the APs of ON-SACs and Ca^2+^ transients in the GCL during cholinergic waves, which suggests that APs enhance not only the postsynaptic ACh-receptor-mediated depolarization-induced Ca^2+^ influx in S-spiking ON-SACs but also presynaptic AP-dependent signal transmission to neighboring ON-SACs. The latter also occurs in cholinergic waves in the INL, as evidenced by how the number of APs in S-spiking ON-SACs is relatively correlated with Ca^2+^ signals in the INL during cholinergic waves. Notably, although the coefficients of determination between the APs and Ca^2+^ transients in both the GCL and INL were significant, their values were relatively low, possibly due to photobleaching of the genetically encoded Ca^2+^ indicator GCaMP6f under fluorescent light (Figure 6B, C). Although two-photon calcium imaging causing less photobleaching is needed to validate our data, our current results suggest that APs modulate the Ca^2+^ wave strength of cholinergic waves. This modulatory role could be necessary for the output of SACs to RGCs because cholinergic waves regulate the development of RGCs and, through RGCs, help to refine eye-specific segregation and retinotopic maps [6–11]. The function of SACs in glutamatergic waves remains unclear. Because they also contain GABA [35,36], SACs could provide GABAergic feedback to bipolar cells and feedforward to RGCs during glutamatergic waves. Through these modulatory roles, the APs of ON-SACs could shape glutamatergic transmission from bipolar cells and RGCs, ultimately refining retinal projections to visual centers of the brain.

## Author Contributions

R.-S.Y. designed and performed the experiments, conducted the data analysis, and wrote the manuscript. X.-L.Y. wrote the manuscript. Y.-M.Z. designed the experiments and wrote the manuscript. D.-Q.Z. designed and oversaw the study and wrote the manuscript. All of the authors approved the final version of the manuscript.

## Funding

This work was supported by the China Scholarship Council (R.-S.Y.), the National Institute of Health Grants (EY022640, D.-Q.Z.), the National Natural Science Foundation of China (81790640, 31872766, and 31571075), the Shanghai Municipal Science and Technology Major Project (No. 2018SHZDZX01; Y.-M.Z.), ZJLab, and the Shenzhen Sanming Project (X.-L.Y.).

## Conflicts of Interest

The authors declare no conflict of interest.

## Notes

### Competing Interest Statement

The authors have declared no competing interest.

